# Intra-Protein Interfaces Control Folding Dynamics and Mechanical Stability in a *De Novo* Designed Repeat Protein

**DOI:** 10.64898/2026.07.31.741961

**Authors:** Melanie Weiß, Anna Lisa Heit, Lukas F. Milles, Johannes Stigler

## Abstract

Protein design has made it possible to generate stable structures with high accuracy, but the principles that determine how designed proteins fold and respond to force remain less understood. Repeat proteins provide a powerful test case because their stability is distributed across modular structural units and nearest-neighbor interfaces rather than concentrated in a single hydrophobic core. Here, we used single-molecule optical tweezers, intramolecular crosslinking and molecular dynamics simulations to dissect the folding landscape of DHR14R, a *de novo* designed helical repeat protein with a deep-learning-redesigned sequence. By introducing site-specific intramolecular crosslinks, we imposed defined boundary conditions on the repeat array and directly manipulated its mechanical folding pathway. DHR14R unfolds reversibly from its mechanically weakest boundary, the N-terminus, through a series of discrete intermediates. Refolding begins with the formation of a three-helix seed rather than a complete two-helix repeat, showing that the structural repeat is not the elementary cooperative folding unit. Preformed terminal interfaces accelerate folding, indicating that seed formation is rate-limiting. Stabilizing both termini eliminates the low-force terminal-fraying pathway and forces disruption of internal interfaces, increasing mechanical stability more than twofold. Together, these results show that designed repeat-protein folding is governed by seed formation, interface propagation, and terminal boundary conditions. They establish intramolecular crosslinking as a strategy for rationally reshaping folding landscapes in designed proteins.

## INTRODUCTION

Protein design has reached a stage where stable three-dimensional structures can be generated with remarkable accuracy. However, designing a structure is not the same as designing a folding landscape. For many applications, including biomaterials [1], molecular devices, and synthetic binding scaffolds [2–4], function depends not only on the final structure but also on how the protein folds, unfolds, and responds to force. Understanding how designed proteins navigate their folding landscapes, and how these landscapes can be rationally redirected, remains a central challenge.

Repeat proteins provide a powerful framework for addressing this problem. Unlike compact globular proteins, whose stability is often organized around a central hydrophobic core, repeat proteins are built from arrays of local structural units coupled by nearest-neighbor interfaces [5, 6]. This architecture creates folding landscapes in which local repeat stability, inter-repeat coupling, and terminal boundary conditions can, in principle, be tuned separately. Helical repeat proteins are especially attractive in this context because they form elongated solenoid or superhelical structures from repeated helix-containing motifs [7, 8]. *De novo* designed helical repeat proteins (DHRs) expand this design space further by providing stable repeat architectures that are not constrained by natural evolution [9].

A defining feature of repeat-protein folding is the importance of inter-repeat interfaces. In many repeat families, the energetic contribution of interfaces between neighboring repeats exceeds the intrinsic stability of isolated repeats. This has been shown for natural repeat proteins such as PR65 [10], designed ankyrin repeat proteins (DARPins) [11, 12], and consensus tetratricopeptide repeat proteins (CTPRs) [13, 14]. Consistent with this interface-dominated architecture, repeat proteins often fold by nucleation and propagation: formation of an initial folded seed or nucleus is rate-limiting, whereas subsequent folding proceeds more rapidly through growth along stabilizing interfaces [15–17]. Terminal repeats are expected to be especially influential because they have fewer stabilizing neighbors and can therefore act as weak points that define folding and unfolding pathways [17, 18].

Most studies of repeat-protein folding have relied on ensemble measurements such as thermal or chemical denaturation, circular dichroism spectroscopy, or stopped-flow kinetics [13, 16, 19–21]. These approaches have established important principles of stability and cooperativity, and targeted mutations have been used to tune repeat-protein stability in ankyrin and TPR systems [22–24]. However, ensemble measurements generally cannot directly resolve the order of intermediates populated by individual molecules, distinguish terminal initiation from internal interface rupture, or determine how defined structural constraints reroute folding under load. Single-molecule force spectroscopy provides a complementary approach because it can resolve discrete folding intermediates and measure folding and unfolding kinetics along a defined mechanical coordinate.

Here, we asked whether the folding landscape of a designed repeat protein can be actively redirected by engineering its terminal boundary conditions. We used single-molecule optical tweezers, site-specific intramolecular crosslinking, and coarse-grained simulations to dissect the folding of DHR14R, a *de novo* designed helical repeat protein with a deep-learning-redesigned sequence. Crosslinks allowed us to stabilize selected terminal repeats, isolate them from force-induced unfolding, and test whether preformed interfaces act as folding seeds. This strategy directly probes whether mechanical unfolding proceeds through terminal fraying or internal interface rupture.

We show that DHR14R unfolds preferentially from its weaker N-terminal boundary through a series of discrete intermediates. Refolding begins with the formation of a three-helix seed rather than a complete two-helix repeat, demonstrating that the structural repeat is not the elementary cooperative folding unit. Stabilizing terminal repeats accelerates folding by bypassing this rate-limiting seed-formation step and, when both termini are crosslinked, reroutes unfolding toward disruption of internal interfaces, strongly increasing mechanical stability. These results show that designed repeat-protein folding is controlled by seed formation, interface propagation, and terminal boundary conditions, and they establish crosslinking as a strategy for rationally reshaping folding landscapes in modular protein architectures.

## RESULTS

### Characterizing single-molecule folding of the *de novo* designed helical repeat protein DHR14R

To characterize the role of inter-repeat interfaces on the folding kinetics, energetics, and mechanical stability, we selected *de novo* designed helical repeat proteins (DHRs) as a model system. DHR14R is a proteinMPNN [25, 26] redesign variant of DHR14 [9], consisting of 160 amino acids in four helix-loop-helix repeats as depicted in fig. 1A. The straight rod-like geometry of DHR14 is achieved by repeating a simple helix-loop-helix pattern with 17 amino acids per helix and three amino acids per loop. Redesigning the protein resulted in structural repeats rather than sequence repeats, mimicking properties of natural repeat proteins. DHR14R showed thermal stability up to 95°C in circular dichroism spectroscopy, which is comparable to the original DHR14 [9] (fig. S1).

**FIG. 1.**
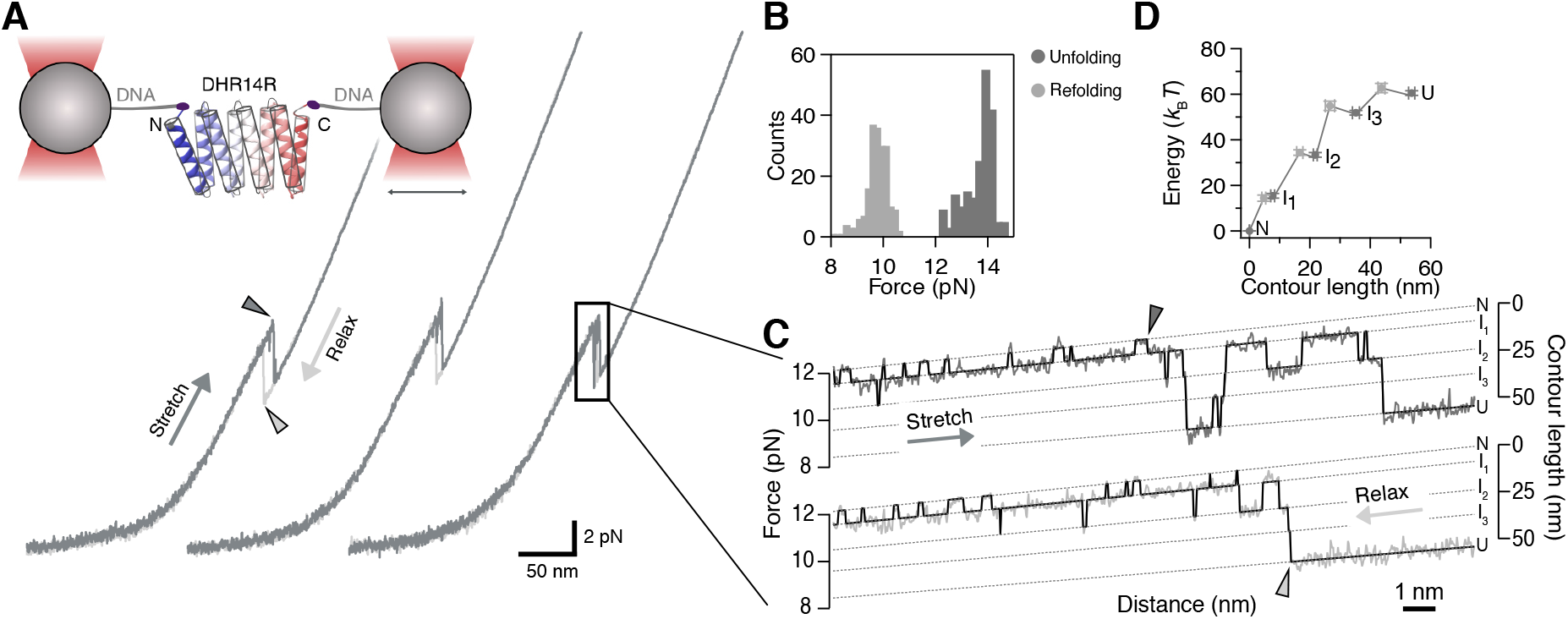
Structure and mechanical properties of the *de novo* designed helical repeat protein DHR14R. (**A**) Top: Scheme of our dual optical tweezers set-up with the optically trapped DHR14R. The structure of the DHR14R construct was predicted by Alphafold2 and is colored from N-terminus (blue) to C-terminus (red). Bottom: Representative force-distance curves (FDCs) of DHR14R pulling at 100 nm*/*s. Darker shades show stretching of the tether leading to protein unfolding, while lighter shades present tether relaxing and protein refolding.**(B)** Histograms of the quantified unfolding (dark) and refolding (light) forces (*n* = 9 molecules with 196 (un)folding events). **(C)** Detailed view of stretching and relaxation curve with fitted hidden Markov model (black). Dashed lines are worm-like chain fits to the native state (N), intermediate states (I_1_–I_3_), and the fully unfolded state (U). Arrowheads indicate the reported unfolding (dark gray) and refolding force (light gray) quantified from the shown FDCs. **(D)** Corresponding interpolated energy landscape of DHR14R. Transition state positions are depicted in light gray. Inter-state barrier heights were estimated using the Arrhenius equation (eq. (S10)). Error bars show statistical errors (SD) from *n* = 9 molecules.

To study the unfolding and folding of DHR14R, we tethered the protein between two optically trapped beads and recorded force-distance curves (FDCs) by performing stretch-relax cycles at a constant velocity of 100 nm*/*s (fig. 1A). We observed reversible unfolding (dark arrowheads) of the protein at 13.5 ± 0.8 pN and refolding (light arrowheads) at 9.6 ± 0.4 pN, on average (fig. 1B). Thus, the protein displays a small hysteresis of a few *k*_B_*T* appearing as the area between unfolding and refolding curves (fig. 1A). We fitted worm-like chain (WLC) models to extract the change in unfolded contour length upon protein unfolding. We found a change in contour length of 53.5 ± 1.1 nm, closely matching the prediction from an Alphafold 2 model [27] of 55.4 nm, indicating that the protein fully unfolds and readily returns to its native structure.

Taking a closer look at the recorded FDCs, we observed hopping between a set of well-defined intermediate states (I_1_–I_3_) between the native (N) and unfolded state (U) (fig. 1C), at positions of 7.7 ± 0.8 nm, 21.8 ± 0.7 nm and 35.3 ± 1.4 nm, respectively and a total difference in free energy upon folding of 58.9 ± 1.1 *k*_B_*T* (fig. 1D). Based on the measured changes in contour length, the N ↔ I_1_ transition is compatible with (un)folding of a single helix, while the I_1_ ↔ I_2_ as well as the I_2_ ↔ I_3_ transition are each compatible with additional (un)folding of two helices, further supported by coarse-grained (CG) simulations (see fig. S2A and video S1).

We previously adopted the use of Ising models [8, 11, 23, 28] for the folding of repeat proteins in single-molecule optical tweezers [14]. We fitted a previously described heteropolymer Ising model [14] to averaged FDCs and found energetic contributions of ≈ 0.5 *k*_B_*T* for the intra-repeat energy Δ*G*_unit_ and ≈ −19 *k*_B_*T* for the inter-repeat energy Δ*G*_nn_ (SI and fig. S3). This indicates that the primary energetic contribution stabilizing DHR14R lies in the interfaces, in line with findings for other repeat proteins [8, 12, 15–17, 29, 30]. Consequently, we expect that unfolding must start from either terminus. In line with this, the contour length change of the *N* ↔ I_1_ transition is compatible with the length expected from the unfolding of one helix from either end (≈ 8 nm). We conclude that DHR14R’s folding behavior mechanistically closely resembles that of other natural and nature-derived repeat proteins.

### Manipulating the folding pathway by stabilizing the termini with intramolecular crosslinks

In mechanical unfolding, the termini are the weakest part. We therefore reasoned that terminal interface stabilization would have substantial impact on mechanical protein stability. To investigate the role of terminal interface stabilization, we introduced intramolecular crosslinks into DHR14R to isolate terminal repeats from force application. To this end, we mutated up to four residues to cysteines located at the beginning of helix 1 and 2 as well as helix 7 and 8, which form intramolecular disulfide bonds under oxidizing conditions and which can resist forces exceeding hundreds of pN [31–33]. This strategy enabled the generation of either a N-terminal crosslink (helix 1 and 2, DHR14R-XN, fig. 2A) or a C-terminal crosslink (helix 7 and 8, DHR14R-XC, fig. 2B), and both simultaneously (DHR14R-XNC, fig. 2C). Consistent with expectations, crosslinking reduced the contour length of unfolding for each variant (38.5 ± 1.5 nm for DHR14R-XN, 35.6 ± 0.8 nm for DHR14R-XC, 24.0 ± 0.9 nm for DHR14R-XNC, fig. S4). The unmodified DHR14R displayed unfolding and refolding via three distinct intermediate states. To further understand the (un)folding pathway of the mutants, we fitted Hidden-Markov models to the FDCs of the crosslinked constructs (fig. 2D–F). DHR14R-XN showed un- and refolding via two intermediate states with changes in contour length of 8.5±0.6 nm for I_1_, and 23.5±1.1 nm for I_2_ (fig. 2D). We observed a similar pattern for DHR14R-XC, unfolding via two intermediate states with length changes of 7.2 ± 0.4 nm for I_1_, and 20.7 ± 0.5 nm for I_2_ (fig. 2E). The measured changes in contour length are compatible with the (un)folding of a single helix (*N* ↔ I_1_), and the (un)folding of a repeat, i.e., two helices (I_1_ ↔ I_2_), further supported by coarse-grained (CG) steered molecular dynamics simulations (fig. S2, video S2 and video S3).

**FIG. 2.**
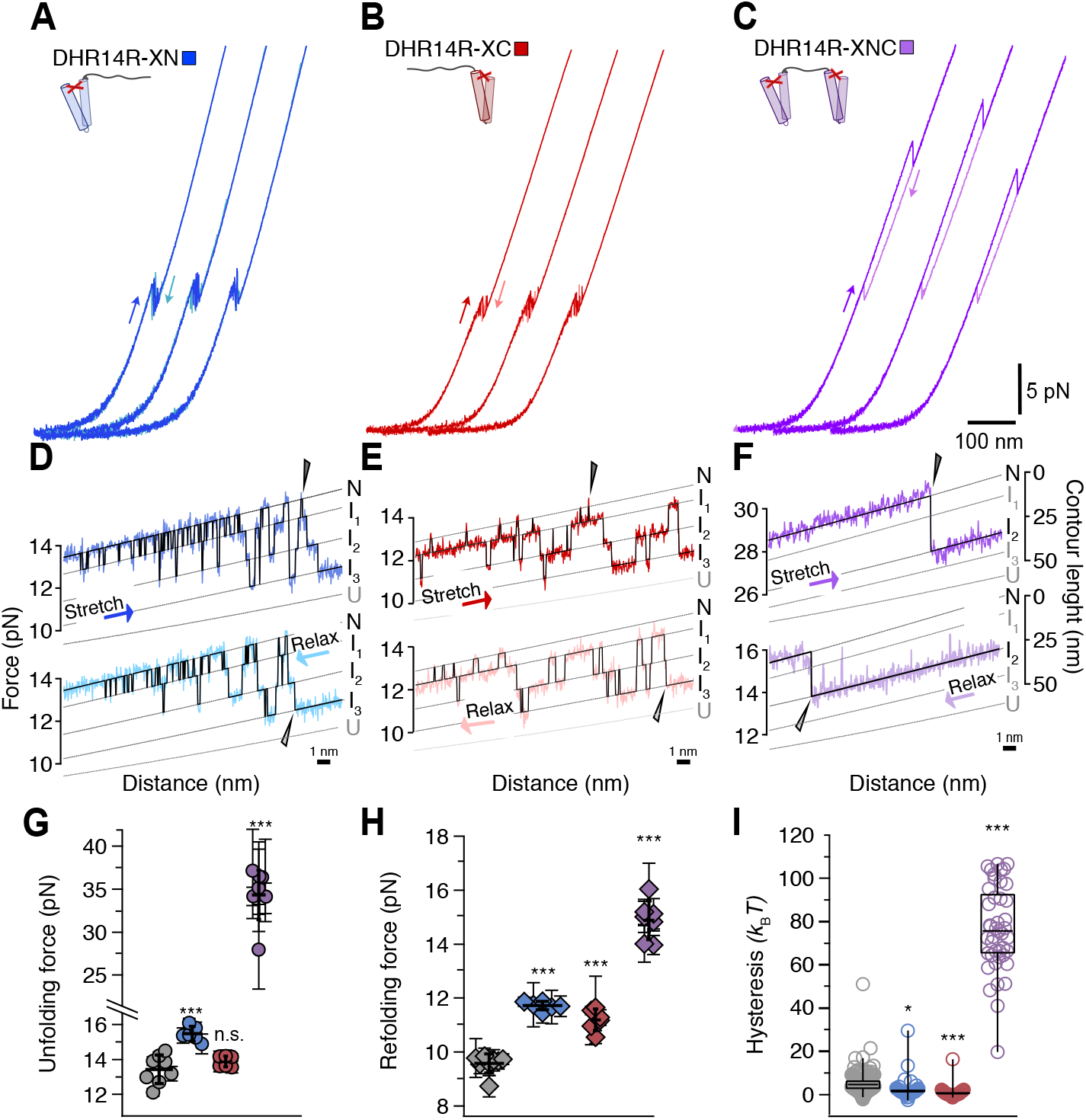
Manipulating the folding pathway of a designed helical repeat protein by intramolecular crosslinks. **(A–C)** Representative FDCs for DHR14R with N-terminal (blue), C-terminal (red), and N- and C-terminal crosslink (purple). Stretch in darker shades, relax in lighter shades. Pulling velocity of 100 nm*/*s. **(D– F)** Zoomed in stretch (dark shades) and relax (light shades) traces with fitted Hidden Markov Model (black trajectories). **(G**,**H)** Quantified unfolding (circles) and refolding (diamonds) forces for all variants. Data points represent the average of 3 to 92 FD cycles for individual molecules. Error bars are standard deviations. **(I)** Boxplots of determined hysteresis between unfolding and refolding curves per construct. One circle represents the quantification of one FD curve. *n* = 6 to 9 molecules per construct. Whiskers show the 2nd and 98th percentiles. Coloring: no crosslink (DHR4R) in gray, N-terminal crosslink (DHR14R-XN) in blue, C-terminal crosslink (DHR14R-XC) in red, N- and C-terminal crosslink (DHR14R-XNC) in purple. Statistical significance was determined using Student’s *t*-tests and is reported relative to DHR14R.

We conclude that the single-crosslinked variants (XN and XC) unfold via the same macroscopic transitions as DHR14R (first a single terminal helix, followed by two-helix packages). In contrast, DHR14R-XNC shows only a single (un)folding transition in experimental data (fig. 2C,F) and CG molecular dynamics simulations (fig. S2B,video S2). The absence of such a transition in DHR14R highlights that an unfolded core with folded terminal repeat is a high-energy intermediate specifically induced by crosslinking both termini, which is not part of the DHR14R folding network.

### DHR14R unfolds from the N-terminus

The pulling traces of the non-crosslinked DHR14R did not allow us to distinguish whether unfolding proceeds via the N-terminus, the C-terminus, or perhaps a mixture of both. We thus compared the variants with single terminal crosslinks, DHR14R-XN and DHR14R-XC, with the WT. While crosslinking the N-terminal repeat (DHR14R-XN) increased the mechanical stability by about 2 pN to 15.5 ± 0.4 pN (*p* = 0.007), the C-terminally crosslinked variant was similar, within error, to the WT (13.9 ± 0.3 pN, *p* = 0.5). We conclude that the N-terminus is the mechanically weakest part and, in the WT, unfolding predominantly proceeds via the N-terminus.

### Crosslinking both termini significantly enhances the mechanical stability of the folded conformation

To compare whether terminal shielding further increases mechanical stability and how it affects the folding pathway and kinetics, we studied DHR14R-XNC, the construct containing crosslinks in both termini (fig. 2C). Strikingly, compared to DHR14R, DHR14R-XNC displayed a substantial increase in mechanical stability to 34.3 ± 3.8 pN, more than double that of the un-crosslinked variant. Interestingly, no intermediate states were populated; instead, DHR14R-XNC unfolded irreversibly in one step (fig. 2F). This absence of intermediates is consistent with the model that breaking internal interfaces is energetically highly disfavored. We conclude that preventing terminal unfolding massively enhances the mechanical stability.

Crosslinking either terminus significantly increased the refolding forces from about 9.6 pN to 11.7 ± 0.4 pN (DHR14R-XN, *p* = 0.00004) or 11.3 ± 0.5 pN (DHR14R-XC, *p* = 0.0002, fig. 2H). Simultaneous crosslinking of both termini (DHR14R-XNC) resulted in an even greater increase of the refolding force to 15.0 ± 0.7 pN. This refolding force increase of 5.5 ± 1.2 pN is comparable to the sum of the force increase for DHR14R-XN and DHR14R-XC (3.9 ± 1.9 pN). Further, we observed a reduction in hysteresis (fig. 2I), quantified as the area between unfolding and refolding curves, for single crosslinked variants, DHR14R-XN (*p* = 0.02) and DHR14R-XC (*p* ≤ 0.0001), compared to the non-crosslinked variant. This suggests increased folding reversibility under load, likely due to enhanced folding kinetics. In contrast, the significant gain in hysteresis of the double-crosslinked DHR14R-XNC construct of 76.9 ± 3.0 *k*_B_*T* compared to 5.1 ± 0.3 *k*_B_*T* for DHR14R (fig. 2I, *p* ≤ 0.0001) suggests that extensive interface stabilization increases the unfolding barrier. As a result, the system becomes more mechanically resistant and farther from equilibrium during constant-velocity force-distance cycling. Together, the results indicate that the introduction of crosslinks prevents the protein from becoming fully denatured at high tension because the applied force must propagate through the crosslink, limiting the parts of the protein that can be unfolded. We thus propose that the crosslinked and tension-isolated termini serve as pre-formed interfaces that facilitate protein folding at higher tension.

### Initial folding seed formation rate-limiting for folding

The introduction of an intramolecular crosslink not only influenced the mechanical properties of DHR14R by guiding the folding/unfolding pathway in a certain way, but it also altered the folding kinetics. Upon introducing a single terminal crosslink, folding/refolding transitions under force occurred more frequently (e.g., compare fig. 2D,E with fig. 1). In addition, faster folding kinetics are reflected by a significant decrease in hysteresis between unfolding and refolding curves for the crosslinked construct compared to the wildtype (fig. 2I). Since the difference between the constructs is the presence or absence of a force-isolated folded repeat, we propose that the formation of a first folded “seed” is rate-limiting. Once a repeat is folded, it serves as a seed for the folding of neighboring repeats by providing a stabilizing interface.

To quantify this effect, we compared the kinetics of the first folding transition in the folding pathway, i.e., the folding/unfolding of the first repeat for WT DHR14R and the folding/unfolding of the first non-crosslinked repeat for DHR14R-XN, DHR14R-XC and DHR14R-XNC (fig. 3). The force-dependent folding (fig. 3A) and unfolding (fig. 3B) kinetics of this first transition are nearly identical for DHR14R-XN, DHR14R-XC and DHR14R-XNC, but different from DHR14R (also see fig. S5). Fitting the data to a two-state model, which includes effects of non-linear tethers ([34], also see methods), allowed inferring the folding rates at zero force *k*_0_, the free energy difference Δ*G* of this transition, and the transition state position Δ*L* (fig. 3D–F). In this model, Δ*L* is defined as the position from the transition state to the unfolded state, on a contour-length scale. Hence, lower values of Δ*L* correspond to a transition state closer to the unfolded state. Consistent with the visual comparison discussed above, the kinetics of the first folding transition for DHR14R-XN (*k*_0_ ≈ 10^6.3^ s^−1^), DHR14R-XC (*k*_0_ ≈ 10^6.5^ s^−1^) and DHR14R-XNC (*k*_0_ ≈ 10^6.5^ s^−1^) are indistinguishable (*p >* 0.25). However, the *k*_0_ rates for all crosslinked mutants differ significantly from the initial formation of a folded unit (*k*_0_ ≈ 10^5.7^ s^−1^, DHR14R data, *p <* 0.035).

**FIG. 3.**
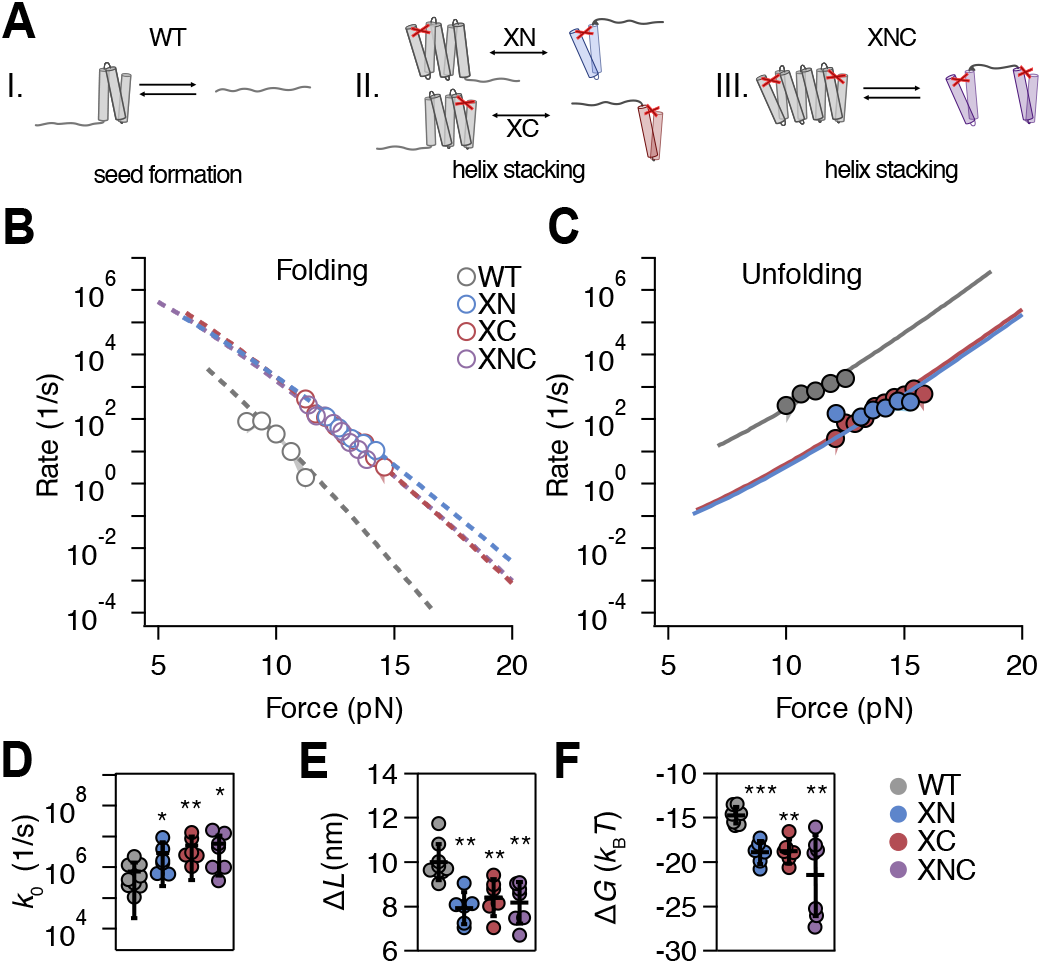
First folding transition kinetics for the DHR14R variants. **(A)** Schematic representation of the first folding transition observed for the different constructs. Scenario I: transition between the fully unfolded state and the folding seed. Scenario II: stacking of neighboring helices to the folded crosslinked repeat for DHR14R-XN (upper scheme in blue) and DHR14R-XC (lower scheme in red). Scenario III: Transition between the native conformation containing two terminal crosslinks and the unfolded core region for DHR14R-XNC. Folding **(C)** Unfolding **(E–F)** Fit parameter. For DHR14R-XNC, analysis was done with 500 nm*/*s stretch-relax cycles since we obtained too few data points to properly fit the chevron plots at 100 nm*/*s. For the other variants 100 nm*/*s, cycles were analyzed.

These kinetic differences suggest that the first folding transition in the single-crosslinked variants is primarily dominated by helix stacking, whereas the WT transition involves initial helix folding. All-atom molecular dynamics (MD) simulations of the isolated N- and C-terminal repeat of DHR14R further suggest that the crosslinked repeat remains weakly folded even if neighboring repeats of the protein are absent (fig. S6). Additionally, the transition state position of DHR14R Δ*L* = 10.0 ± 0.8 nm differs significantly (*p <* 0.005) from that of all crosslinked variants (Δ*L* ≈ 8 nm), indicating that different macroscopic transitions are observed.

The observation that Δ*G* of the non-crosslinked DHR14R differs from that of the crosslinked variants -XN and -XC (fig. 3E) is consistent with the formation of more interfaces when forming the three-helix containing “seed” in DHR14R, vs the stacking of two helices in DHR14R-XN or DHR14R-XC. To verify whether three helices are sufficient for folding seed formation, we generated a truncated construct DHR14R-3H containing only the three C-terminal helices of DHR14R. Consistent with expectations, DHR14R-3H perfectly reproduced the kinetics and change in contour length of the first folding transition of DHR14R within errors (fig. S7). Further, in CG simulations of WT DHR14R unfolding, the last unfolding transition corresponds to the (un)folding of three helices (fig. S2A, video S1). These results clearly indicate that the folding seed of DHR14R is formed by a 3-helix bundle rather than by a single repeat.

Together, these results suggest that the formation of a folding seed is the rate-limiting step in DHR14R protein folding and that the presence of a folding seed increases the folding kinetics. These findings further highlight the crucial role of seed formation in repeat protein folding by contributing folding interfaces.

### Changes in folding properties exclusively originate from introduced crosslinks

To verify whether the observed changes in mechanical properties and folding kinetics are exclusively due to the presence of a folding seed rather than the introduced mutations, we performed control experiments for the disulfide crosslinked constructs under reducing conditions. All crosslinked variants were measured in buffer containing 1 mM dithiothreitol (DTT) to reduce the disulfide bonds. Indeed, reduction of the disulfide bonds reproduced the WT behavior for all constructs (fig. S8). Serendipitously, we observed the stepwise reduction of both disulfide bonds in a single DHR14R-XNC molecule in real time during pulling and relaxing cycles (fig. 4), as evidenced by a step-wise lengthening of the polypeptide chain. A direct comparison of the FD curves of this molecule with both crosslinks intact (1., 2.) with the curve of one reduced crosslink (3.–5.) shows not only chain lengthening but also an increase in folding/unfolding kinetics as expected for the reversal of one crosslink. Consistent with expectations, reversal of the second terminal crosslink (6.–8.) reproduced the unfolding pattern and kinetics of DHR14R (additional FDCs in fig. S8D). We conclude that the observed effects are attributable to the formation of crosslinks, rather than cysteine replacement.

**FIG. 4.**
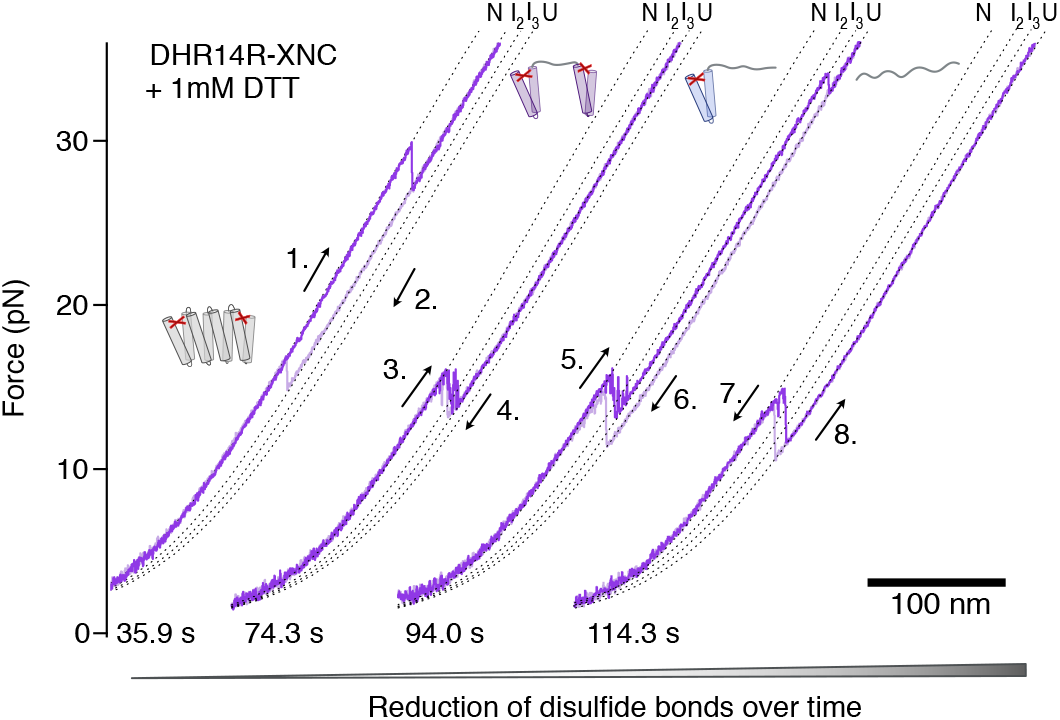
Gradual crosslink reduction in DHR14R-XNC with 1 mM DTT. Shown FDCs are consecutive observations for the same molecule, and the chemical reduction of disulfide bridges was observed in real time. The data were measured at a pulling velocity of 100 nm*/*s and span about 80 s, as indicated by the time-stamps. Cartoons depict the unfolded state, depending on the disulfide bonds formed/reduced.

## DISCUSSION

### Designed repeat proteins as models for modular folding

Repeat proteins differ from compact globular domains because their stability is distributed across a linear array of structural units and depends strongly on nearest-neighbor interactions [5, 6]. This modular architecture makes them useful systems for dissecting how local repeat stability, inter-repeat interfaces, and terminal boundary conditions shape folding landscapes [11, 15, 16, 28, 29]. Here, we used the *de novo* designed helical repeat protein DHR14R to define how these parameters control folding, unfolding, and mechanical stability at the single-molecule level.

DHR14R behaves as a repeat protein in the physical sense. It unfolds and refolds through discrete intermediates whose contour-length changes correspond to the sequential loss and gain of helical units. Ising-model analysis indicates that the dominant energetic contribution arises from inter-repeat interfaces rather than from the intrinsic stability of individual repeats, consistent with natural and consensus repeat proteins [8, 11, 14, 16, 28]. Thus, designed repeat proteins can reproduce central physical principles of natural solenoidal proteins while providing a controllable platform for perturbing them [9, 25, 35, 36].

### Terminal asymmetry defines the unfolding pathway

The termini of repeat proteins are expected to be mechanically vulnerable because terminal repeats have fewer stabilizing neighbors [5, 6]. Our crosslinking experiments directly show that the two termini of DHR14R are not mechanically equivalent. Stabilizing the N-terminal repeat increased the unfolding force, whereas stabilizing the C-terminal repeat had little effect. Thus, DHR14R unfolds predominantly from the N-terminus, identifying this region as the dominant mechanical initiation site.

This directionality is not necessarily universal. Ankyrin repeat proteins have been reported to unfold predominantly from the N-terminus and fold from the C-terminus [22, 37], whereas simulations of Gankyrin suggested C-terminal unfolding initiation [17]. Terminal unfolding, therefore, likely reflects the local energetic architecture of a given repeat array rather than an intrinsic preference for one end. Furthermore, the geometry of a repeat itself can influence directional unfolding as seen for CTPRs [14]. However, DHR14R displays a straight, rod-like geometry and therefore the geometry effect of directional unfolding is unlikely. We speculate that, in our case, the observed predominant unfolding from the N-terminus is rather based on the amino acid sequence composition than on the geometry since DHR14R does not contain sequence repeats. When comparing the H-bonds in our construct, we found that the C-terminal repeat globally forms eight more H-bonds compared to the N-terminal repeat. This dissimilarity in the number of formed H-bonds could at least partially explain the stability difference of the termini. Additionally, the total free energy for XN and XC differs by 9 *k*_B_*T*, strongly suggesting cap asymmetry. In DHR14R, the different effects of N- and C-terminal crosslinking indicate that local sequence, packing, or interface energetics make the N-terminal boundary weaker. Our data extend earlier work on tuning repeat-protein stability by local mutations or cap modifications [16, 23, 38, 39] to mechanical unfolding under load.

### Crosslinks reroute the mechanical folding landscape

The double-crosslinked construct provides the strongest evidence that terminal structure controls the mechanical pathway. When both terminal repeats are protected, DHR14R no longer unfolds through the same low-force, stepwise pathway. Instead, it unfolds in a single high-force transition in which the core is disrupted while the terminal repeats remain structured. This more than twofold increase in unfolding force shows that mechanical stability is determined not only by native-state stability, but also by the topology of the force-bearing pathway.

This agrees with single-molecule studies showing that pulling geometry and unfolding topology strongly affect mechanical resistance [40–43]. In DHR14R, the relevant topology is the accessibility of weak terminal units and the propagation of interface rupture along the repeat array. Terminal crosslinks therefore act as engineered boundary conditions: they remove the low-barrier terminal-fraying route and force unfolding through internal interface disruption.

The disulfide-reduction experiments support this interpretation by showing that stepwise reduction restores longer contour lengths and DHR14R-like behavior, indicating that the altered mechanics arise from the crosslinks rather than from the cysteine substitutions. This is consistent with the high mechanical robustness of disulfide bonds [31–33].

### Folding proceeds by seed formation and interface propagation

The folding data indicate that a folded terminal structure can act as a kinetic seed. Single terminal crosslinks increase refolding forces and reduce hysteresis, consistent with faster, more reversible folding under load. Because a crosslinked terminal repeat likely remains structured as observed in all-atom MD simulations (fig. S6), it provides a preformed interface onto which neighboring helices can assemble. Force-dependent kinetic analysis supports this model: the first folding transition in the crosslinked variants is faster than the initial folding transition of unmodified DHR14R and has a distinct transition-state position. Thus, the slow step in DHR14R folding is the formation of the first productive folded unit, not propagation along an existing interface.

Contour length information of DHR14R, together with data from DHR14R-3H and CG simulations, shows that this initial folding seed is a three-helix bundle rather than a complete two-helix repeat. Under load, DHR14R therefore does not fold by a simple repeat-by-repeat mechanism. Instead, folding proceeds by nucleation and propagation: a minimal multi-helix seed forms first, after which the folded structure grows through stabilizing inter-repeat interfaces. Similar departures from strict repeat-by-repeat folding have been observed in other repeat proteins [13, 14], supporting the view that the structural repeat is not necessarily the elementary cooperative folding unit. Seed formation has also been proposed as rate-limiting in other repeat-protein systems [16, 19].

### DHR14R combines high stability with fast folding

DHRs and other designed proteins are often highly thermostable [9, 19, 36, 38], but less is known about their mechanical stability and folding kinetics. The DHR14R three-helix seed folds on a microsecond timescale (*τ* = 1*/k*_0_ ∼ 1 to 2 µs, comparable to fast-folding helical proteins such as HP35 and to other *de novo* three-helix bundles [44, 45]. These rates are physically plausible relative to proposed folding speed limits [46]. Thus, designed repeat proteins can combine high thermal stability, substantial mechanical stability and efficient folding, provided that their landscapes contain accessible nucleation routes with limited rate-limiting metastable intermediates [47].

### Implications for protein engineering

Natural repeat proteins function as interaction platforms, spacers, elastic elements, and regulatory scaffolds [28, 48, 49]. Many likely experience directional forces during molecular recognition, remodeling, transport, degradation, or mechanical coupling. Our results suggest that modest changes at repeat-protein termini can strongly affect mechanical response by suppressing or promoting terminal fraying.

The crosslinking strategy also provides a route for engineering repeat proteins with programmable mechanical properties. A single terminal crosslink can increase mechanical stability and accelerate refolding, whereas double terminal crosslinking suppresses low-force intermediates and reroutes unfolding. Accordingly, repeat proteins show non-additive mechanical properties. Similar principles may be useful for designing molecular tension sensors, force-bearing linkers, or nanoswitches [50, 51]. For intracellular applications, however, disulfide bonds would likely need to be replaced by more robust covalent constraints, such as autocatalytic isopeptide bonds [52, 53].

## Conclusion

In summary, DHR14R folds through a rate-limiting three-helix seed and then propagates structure through stabilizing inter-repeat interfaces. Mechanical unfolding normally initiates at the weaker N-terminal boundary, but this pathway can be rerouted by stabilizing terminal repeats. By converting weak termini into protected folding seeds, crosslinks accelerate refolding and, when placed at both ends, strongly increase mechanical stability by forcing disruption of internal interfaces. These results establish designed helical repeat proteins as tractable models for modular protein folding and show that mechanical folding landscapes can be reshaped by engineering repeat-protein boundary conditions.

## MATERIALS AND METHODS

### Protein construct design

The designed protein studied here is based on the DHR14 backbone (RCSB: 5CWH) designed by Brunette et al. (2015) [9]. The amino-acid sequence was generated with protein MPNN [25, 26]. For intramolecular crosslinking, cysteines were site-specifically introduced (amino acid 18 and 54 in the N-terminal repeat and amino acid 137 and 176 in the C-terminal repeat, respectively) via Gibson assembly to form disulfide bonds in oxidizing conditions. Alphafold2 [27] was used to verify that the mutations do not change the backbone structure. All proteins contained one N- and one C-terminal ybbR tag [54] for DNA handle attachment [55], as well as a C-terminal SNAC cleavage tag[56], followed by a His-tag for affinity chromatography purification.

### Protein expression and purification

All proteins were expressed in *E. coli* strain BL21(DE3). Cultures were grown at 37 ^◦^C in lysogeny broth (LB) media. Protein expression was induced with 0.1 mM Isopropyl-beta-D-thiogalactopyranosid (IPTG) after an optical density of 0.6 was reached. Cells were harvested after three hours of growth at 37 ^◦^C. Cell lysis was done by sonification and addition of 0.1 mg*/*mL lysozyme. Proteins were purified via His-tag affinity purification using Ni-NTA resin, followed by size exclusion (Superdex 200 cytica column) in PBS (10 mM Na_2_HPO_4_, 1.8 mM KH_2_PO_4_, 137 mM NaCl, 2.7 mM KCl) to isolate monomeric proteins.

### Single-molecule measurements

Proteins of interest were tethered between two optically trapped beads [57, 58] (silica beads, anti-digoxigenin or streptavidin coated, 1 µm diameter) as previously described using N- and C-terminal ybbR tags of the protein to enzymatically attach Coenzyme A (CoA) modified DNA oligos (Biomers), which later hybridized to 545 bp DNA handles [14, 55, 59]. Oligo attachment was done by incubating protein of interest with 2x excess of sfp and 2.5x excess of CoA-oligo supplemented with 10 mM MgCl_2_ for two hours at room temperature. Size-exclusion chromatography was performed to confirm DNA-protein conjugation and isolate proteins with N- and C-terminal conjugated DNA. DNA handles were generated by polymerase chain reaction using dual-bio or dual-dig forward primer and a backward primer containing an abasic site to generate a 34-base single-stranded overhang for hybridization to the CoA-oligo, as previously described [59]. A dual optical trap (Lumicks, C-trap) with custom-built measurement chambers was used for trapping experiments. All trapping experiments were performed at 298 K in PBS (1 mM DTT added for reducing conditions), supplemented with oxygen scavenger system (0.8 % w/v glucose, 90 U*/*mL glucose oxidase, and 1700 U*/*mL catalase). Prior to measurements, chambers were passivated for 5 to 10 min with 10 mg*/*mL bovine serum albumin (BSA), followed by three to four washing steps with PBS and sample application. Force distance measurements were performed with pulling velocities of 100 nm*/*s and trap stiffness around 0.35 pN*/*nm, unless indicated otherwise.

### Data analysis

Data analysis was done using custom-written Igor Pro (wave metrics) scripts. Data was acquired at 78.125 kHz. The DNA tether response was fitted using an extensible Worm-like chain model (eWLC), and (partially) unfolded protein states were fitted to WLC models. Kinetics were extracted by fitting a multi-state Hidden Markov Model (HMM), implemented using force-dependent transition rate constants and Viterbi path optimization. Prior to HMM analysis, the raw data was downsampled by a factor of 20 for computational performance. Unfolding and refolding forces were defined as the force where the fitted trajectory last leaves the fit of the native state during unfolding, or first leaves the fit of the unfolded state, respectively. Hysteresis between unfolding and refolding was defined as the area between fitted curves to the stretch and relax curves. Force-dependent folding kinetics were fitted to a two-state implementation of a model that describes the force-dependent energetics of linker stretching [34]. All error bars represent standard deviations. Statistical significance was determined using Student’s *t*-test. Significance levels are as follows: *p* ≥ 0.05 was set as not significant (n.s.), *p <* 0.05 is indicated with *, *p <* 0.01 **, and *p <* 0.001 ***.

## AUTHOR CONTRIBUTIONS

Investigation: MW; methodology: MW; data analysis: MW, JS; Simulations: JS, ALH; protein construct design: LFM; data visualization: MW; programming: JS; writing: MW, JS; supervision: JS.

## ACKNOWLEDGMENTS

We thank Marvin Freitag for initial help with setting up and optimizing the single-molecule experiments, Stephan Uebel for access to the CD spectrometer, Joelle Deplazes-Lauber and Jeanny Probst for helping with cloning and technical support. We further thank the whole Stigler lab for thoughtful discussion and scientific input. MW acknowledges support from the Graduate School of Quantitative and Molecular Biology (QMB), Munich, and the LMU Center of Nanoscience (CeNs). This work was funded by Deutsche Forschungsgemeinschaft under grant 527129816.

