## Supplementary Information for "Intra-Protein Interfaces Control Folding Dynamics and Mechanical Stability in a *De Novo* Designed Repeat Protein"

### I. PROTEIN SEQUENCES

#### A. DHR14R wildtype with tags

SDSLEFIASKLAGSSGEEELKKKVEELAEKAKKEKDPEEIIKLAKELVELAKLTES
EEIVKEIVRELAEIAKKEKDPETIIKIAKLLVELAKLTESEEIVKEIVRELAEIAKKEKD
PETIIKIAKLLLELAELTENSEIIKEIKRELEEIAKKEKDPETIEYIKKLLEKLKELSKES
GGSGDSLEFIASKLAGGSHHWGSHHHHHH

#### B. DHR14R-XN with tags

SDSLEFIASKLAGSSGECELKKKVEELAEKAKKEKDPEEIIKLAKELVELAKLCES
EEIVKEIVRELAEIAKKEKDPETIIKIAKLLVELAKLTESEEIVKEIVRELAEIAKKEKD
PETIIKIAKLLLELAELTENSEIIKEIKRELEEIAKKEKDPETIEYIKKLLEKLKELSKES
GGSGDSLEFIASKLAGGSHHWGSHHHHHH

#### C. DHR14R-XC with tags

SDSLEFIASKLAGSSGEEELKKKVEELAEKAKKEKDPEEIIKLAKELVELAKLTES
EEIVKEIVRELAEIAKKEKDPETIIKIAKLLVELAKLTESEEIVKEIVRELAEIAKKEKD
PETIIKIAKLLLELAELTENCEIIKEIKRELEEIAKKEKDPETIEYIKKLLEKLKELCKES
GGSGDSLEFIASKLAGGSHHWGSHHHHHH

#### D. DHR14R-XNC with tags

SDSLEFIASKLAGSSGECELKKKVEELAEKAKKEKDPEEIIKLAKELVELAKLCES
EEIVKEIVRELAEIAKKEKDPETIIKIAKLLVELAKLTESEEIVKEIVRELAEIAKKEKD
PETIIKIAKLLLELAELTENCEIIKEIKRELEEIAKKEKDPETIEYIKKLLEKLKELCKES
GGSGDSLEFIASKLAGGSHHWGSHHHHHH

#### E. Original DHR14 wildtype with tags

SDSLEFIASKLAGSSGSEEVNERVKQLAEKAKEATDKEEVIEIVKELAELAKQSTD
SELVNEIVKQLAEVAKEATDKELVIYIVKILAELAKQSTDSELVNEIVKQLAEVAKEATD
KELVIYIVKILAELAKQSTDSELVNEIVKQLEEVAKEATDKELVEHIEKILEELKKQSTD
SGGSGDSLEFIASKLAGGSHHWGSHHHHHH

#### F. DHR14R N-terminus for MD simulations

SDSLEFIASKLAGSSGEGSSGEEELKKKVEELAEKAKKEKDPEEIIKLAKELVELAKLTES

##### G. DHR14R C-terminus for MD simulations

TENSEIIKEIKRELEEIAKKEKDPETIEYIKKLLEKLKELSKESGGSGDSLEFIASKLA

##### H. DHR14R crosslinked N-terminus for MD simulations

SDSLEFIASKLAGSSGECELKKKVEELAEKAKKEKDPEEIIKLAKELVELAKLCES

##### I. DHR14R crosslinked C-terminus for MD simulations

TENCEIIKEIKRELEEIAKKEKDPETIEYIKKLLEKLKELCKESGGSGDSLEFIASKLA

YbbR tags colored in purple, His-tag in gray, amino acids that were mutated to cysteines are indicated in red.

#### II. METHODS

##### A. Data analysis

###### 1. *Polymer models*

Elastic responses were modeled as an extensible worm-like chain (eWLC) model [1] describing the response of the DNA linkers

$$F(\xi_{\mathrm{D}}) = \frac{k_{\mathrm{B}}T}{p_{\mathrm{D}}} \left( \frac{1}{4} \left( 1 - \frac{\xi_{\mathrm{D}}}{L_{\mathrm{D}}} + \frac{F}{K} \right)^{-2} - \frac{1}{4} + \frac{\xi_{\mathrm{D}}}{L_{\mathrm{D}}} - \frac{F}{K} \right) \tag{S1}$$

in series with a worm-like chain model [2] describing unfolded polypeptide

$$F(\xi_{\mathrm{p}}) = \frac{k_{\mathrm{B}}T}{p_{\mathrm{p}}} \left( \frac{1}{4} \left( 1 - \frac{\xi_{\mathrm{p}}}{L_{\mathrm{p}}} \right)^{-2} - \frac{1}{4} + \frac{\xi_{\mathrm{p}}}{L_{\mathrm{p}}} \right) \tag{S2}$$

with DNA extension $\xi_{\mathrm{D}}$, persistence length $p_{\mathrm{D}}$, contour length $L_{\mathrm{D}}$, and stretch modulus $K$, as well as protein extension $\xi_{\mathrm{p}}$, protein persistence length $p_{\mathrm{p}} = 0.7\,\mathrm{nm}$, and protein contour length $L_{\mathrm{p}}$. Typical values were $K \sim 1000\,\mathrm{pN}$, $L_{\mathrm{D}} \sim 370\,\mathrm{nm}$ and $p_{\mathrm{D}} \sim 10$ to $25\,\mathrm{nm}$.

Thus, force-distance curves (FDCs) $F(d)$ were modeled by inverting

$$d(F) = \xi_{\mathrm{D}}(F) + \xi_{\mathrm{p}}(F) + F/k_c, \tag{S3}$$

where $k_c = 1/(1/k_1 + 1/k_2)$ is the combined trap stiffness of the two optical traps [3].

The mechanical energy needed to stretch such a construct, consisting of DNA, unfolded polypeptide, and traps up to an inter-trap distance $d$ is

$$G^{\mathrm{mech}}(d, L) = \int_0^d F(d')\,\mathrm{d}d', \tag{S4}$$

where $L = L_{\mathrm{p}}$ describes the unfolded contour length of the polypeptide chain.

*2. Transition rate models*

For a protein with $N$ states, the interconversion between states is described by the tension-dependent transition rate matrix $k_{ij}(d)$, describing the rate for transitioning from state $i$ to state $j$. If $i \to j$ describes a folding transition, the rate was modeled as being dependent on the energy required to reduce the contour length of the unfolded peptide from its original value $L_i$ to that of the transition state $L_i^\ddagger = L_i - \Delta L_i$ [4]. Here, the model parameter $\Delta L_i$ describes the distance of the transition state between $i$ and $j$, measured from $i$. Thus, to fulfill detailed balance, the transition rate matrix is given by

$$k_{ij}(d) = \begin{cases} k_{ij}^0 \exp \frac{-\Delta G_{i\ddagger}^{\text{mech}}(d)}{k_\text{B}T} & \text{if } i \to j \text{ is a folding transition} \\ k_{ji}(d) \exp \frac{G_j^0 - G_i^0}{k_\text{B}T} & \text{if } i \to j \text{ is an unfolding transition} \end{cases}, \tag{S5}$$

where $\Delta G_{ij}^{\text{mech}}(d) = G^{\text{mech}}(d, L_j) - G^{\text{mech}}(d, L_i)$ and $\Delta G_{i\ddagger}^{\text{mech}}(d) = G^{\text{mech}}(d, L_i^\ddagger) - G^{\text{mech}}(d, L_i)$.

The transition rate matrix $k_{ij}(d)$ is parameterized by two classes of parameters: fixed structural parameters determined a priori (DNA parameters $p_\text{D}$, $L_\text{D}$, $K$) and unfolded polypeptide contour lengths $L_i, i = 1 \ldots N$), and free kinetic and thermodynamic parameters initialized as a plausible guesses ($k_{ij}^0$, $\Delta L_i$, $G_i^0$, $i, j = 1 \ldots N$, $i \neq j$), which are further optimized.

*3. State calling and parameter optimization*

The transition rate matrix (eq. (S5)) is translated into a discrete time Markov transition probability matrix

$$T_{ij}(d) = 1 - \exp(-k_{ij}(d) \cdot \delta) \tag{S6}$$

with sampling rate $\delta$. A variant of the Viterbi algorithm, previously adapted for state calling in constant-trap-separation passive-model trajectories [5], is then used to determine the likeliest sequence of states for an experimental noisy FDC. The distance-dependent emission probabilities, i.e., the expected probability distributions for producing a particular force given the protein's state, were obtained from polymer models.

During optimization, the Viterbi likelihood is maximized while the free parameters (see above) are adjusted.

*4. Extraction of apparent folding rates and energies from constant velocity force-distance curves*

Apparent force-dependent transition rates were estimated from Viterbi fits to FD traces using by binned counting of transition events via

$$k(F) = \frac{N(F)}{T(F)}, \tag{S7}$$

where $N(F)$ is the number of transitions and $T(F)$ is the cumulative time spent in the bin at force $F$ [6].

Accordingly, the apparent force-dependent folding rate $k_\text{fold}$ can be describes by

$$k_\text{fold}(F) = k_0 \exp\left(-\frac{G_\text{mech}^\ddagger(F)}{k_\text{B}T}\right). \tag{S8}$$

where $G^{\ddagger}_{\text{mech}}$ is the mechanical free energy difference between the unfolded state and the transition state.

The equilibrium free energy difference $\Delta G_0$ between the folded and the unfolded state is based on the detailed balance principle defined as

$$\Delta G_0 = k_{\text{B}}T \ln \frac{k_{\text{fold}}}{k_{\text{unfold}}}. \tag{S9}$$

Free energy landscapes were reconstituted with the obtained changes in free energy between states and the corresponding fitted contour length. The barrier height was estimated using the Arrhenius equation

$$\frac{\Delta G^{\ddagger}}{k_{\text{B}}T} = \ln \frac{k_0}{A}, \tag{S10}$$

where we assumed $A = 1.2 \times 10^4\,\text{s}^{-1}$ [3, 7].

##### 5. Heteropolymer Ising model

Ising models are widely used in statistical mechanics to describe systems with bistable units that transition between two states, 0 and 1. This framework can be applied to the microscopic states of an $N$-unit protein, where each individual unit is either unfolded (0) or folded (1). Consequently, the entire protein can adopt $2^N$ possible configurations. Here, we applied a simplified Ising model using the zipper approximation [8], in which unfolding starts from the termini. Consequently, configurations with internal repeats unfolded, while neighboring repeats are folded (e.g. 1011) are not allowed.

To extract the contribution of the intrinsic energy $G_{\text{unit}}$ and the interfacial energy $G_{\text{nn}}$ of a repeat, we applied a previously described mechanical Ising model [8] to averaged force-extension curves.

In brief, the model assumes that $G_{\text{int}}$ is dependent on the number of unfolded repeats $n$ according to

$$G_{\text{int}} = nG_{\text{unit}} + (n-1)\,G_{\text{nn}} + G_{\text{caps}} \tag{S11}$$

$G_{\text{caps}}$ is an energetic term accounting for an increased hydrophilic surface due to solvent exposure of the terminal repeats, which shield the hydrophobic core from surrounding solvent.

A DHR helix-loop-helix repeat was modeled containing a type A and a type B helix. Accordingly, a heteropolymer Ising model was anticipated to best describe the DHR14R energy contributions via $\Delta G_{\text{unit}} = nG_{\text{A}} + nG_{\text{B}} + (n-1)\,G_{\text{AB}}$, with $\Delta G_{\text{nn}} = (n-1)G_{\text{AA}} + nG_{\text{BB}} + (n-1)\,G_{\text{BA}}$. Here, $G_{\text{A}}$ is the intrinsic energy of a type A helix, $G_{\text{B}}$ is the intrinsic energy of a type B helix, and $G_{\text{AB}}$ is their coupling energy. Accordingly, $G_{\text{AA}}$, $G_{\text{BB}}$, and $G_{\text{BA}}$ are coupling energies of the respective helices in neighboring repeats. Due to amino acid sequence differences of terminal repeats and their solvent exposure, stabilizing capping energies for the N-terminal cap ($G_{\text{N-cap}}$) as well as for the C-terminal cap ($G_{\text{C-cap}}$) were added to the model as $G_{\text{caps}}$ in eq. (S11).

#### B. Simulations

##### 1. Coarse-grained simulations

Cafemol 3.2.1 [9] was used to simulate constant-velocity pulling and relaxing experiments for all DHR14R variants. Simulations were performed using AlphaFold2 structure predictions of all variants, containing N- and C-terminal ybbR tags. Pulling residues were selected as the serines in the ybbR tags of the protein, where a CoA oligo is conjugated. Constant temperature simulations were performed at 298 K with an ionic strength of 150 mM NaCl. All simulations were run in triplicate with a speed of $0.05 \times 10^{-3}$ and $k = 1 \times 10^{-3}$. As an energy function, the AICG2+ model with flexible local potential was used.

##### 2. *Molecular dynamics simulations*

All-atom molecular dynamics (MD) simulations were performed to simulate the intrinsic stability of the N-terminal and C-terminal repeat of DHR14R in isolation and the absence of applied force. Simulations were carried out using AlphaFold2 structure predictions of the isolated N-terminal and C-terminal repeats of DHR14R, including a neighboring ybbR tag, in triplicate using NAMD 3.0 [10]. To investigate the effect of crosslinks on the stability of the terminal repeats, WT terminal repeats (containing no cysteines) as well as the disulfide-bond crosslinked repeats were simulated. The DISU patch in psfgen was used to parameterize a covalent linkage between cysteine residues. System preparation was performed using VMD (version 1.9.3) [11] and the CHARMM36 [12] force field. The simulated constructs were solvated in a TIP3P water box to allow free movement of the protein and ionized to a 150 mM NaCl concentration using the autoionize plugin in VMD [11]. Energy minimization was performed using NAMD 3.0 [10] over 10,000 steps, followed by equilibration in two sequential phases: 200 ps under NVT conditions (canonical ensemble) and 2 ns under NPT conditions (isothermal-isobaric ensemble). Both phases were conducted at 298 K and 1 atm using a Langevin thermostat and a Langevin barostat. Hydrogen mass repartitioning (HMR) [13] was applied in VMD, redistributing mass from bonded heavy atoms to hydrogen atoms (∼3 amu). This enabled a stable integration timestep of 4 fs, substantially reducing computational cost without compromising simulation accuracy.

Each construct was simulated for 5 µs, divided into consecutive 100 ns segments. To improve the performance, the GPU-accelerated integration was enabled via the CUDASOAintegrate option. Long-range electrostatics were calculated using the Particle Mesh Ewald (PME) method with a grid spacing of 1.2 Å and a real-space cutoff of 12 Å, with a switching function applied from 10 Å. To assess thermal stability and denature secondary structures, additional simulations were performed at 373 K.

Trajectory analysis, including root-mean-square deviation (RMSD) and root-mean-square fluctuation (RMSF) calculations and structural visualization, was performed using VMD (version 1.9.3). RMSD values were analyzed for each terminal repeat, excluding the ybbr tag to compare the intrinsic stability of the N- and C-terminal repeat in the absence and presence of a crosslink. Accordingly, residues 2 to 42 were selected for RMSD/ RMSF calculation of the N-terminal repeat and residues 15 to 55 for the C-terminal repeat, respectively, including backbone atoms. RMSF mean values per construct were set as the mean of the respective RMSF of those 40 repeat residues. Additionally, RMSF scores were recorded for all residues, including the ybbR tag. We set the equilibrated structure after 0.05 µs (frame 10) as the reference structure for all RMSD calculations.

#### C. Thermal stability

Thermal stability was determined by observing the change in circular dichroism (CD) signal at 222 nm upon heating up the protein from 20 °C to 95 °C with 1 °C per minute in PBS. For all protein variants, typical alpha helical CD profiles with minima at 208 nm and 222 nm were still observed at 95°C, indicating that the proteins remained properly folded. This high thermal stability is consistent with the original DHR14 [14] and with other DHRs and designed proteins. Since most designed proteins are highly stable, high temperature and high guanidine (GdHCl) concentrations alone do not lead to complete unfolding of the structures. Thus, a combination of temperature and GdHCl was needed. Accordingly, melting experiments were performed in PBS supplied with 5 M GdHCl as already done by Wetzel et al. (2008) [15]. CD measurements were performed on a Jasco J-715 spectrometer with a Peltier temperature controller using a 1 mm pathlength quartz cell. However, even when performing thermal melting in 5 M GdHCl, the DHR14R constructs still remained folded.

#### III. SUPPLEMENTARY FIGURES

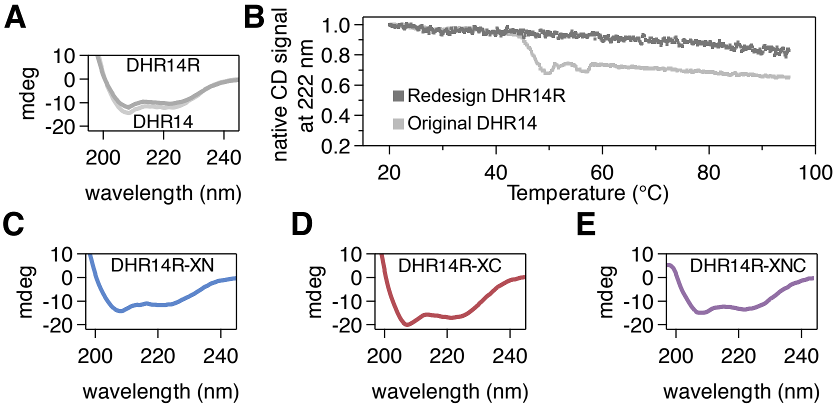

FIG. S1. **(A)** CD spectra for the original DHR14 (lighter gray, lower spectrum) and the redesigned DHR14R (darker gray, upper spectrum) from 197 nm to 245 nm at 20 °C in PBS. **(B)** Melting curves for the original DHR14 (lighter gray) and the redesigned DHR14R (darker gray) in PBS supplied with 5 M GdHCl. The change in CD signal was monitored at 222 nm and the fraction of native CD signal is plotted against the temperature gradient. CD spectra for the crosslinked variants DHR14R-XN **(B)**, DHR14R-XC **(C)**, DHR14R-XNC **(D)** respectively. All CD spectra show a similar characteristic alpha-helical spectrum. No thermal melting was observed for all constructs when applying a temperature gradient from 20 °C to 95 °C in PBS as well as in PBS supplied with 5 M GdHCl for the DHR14R variants.

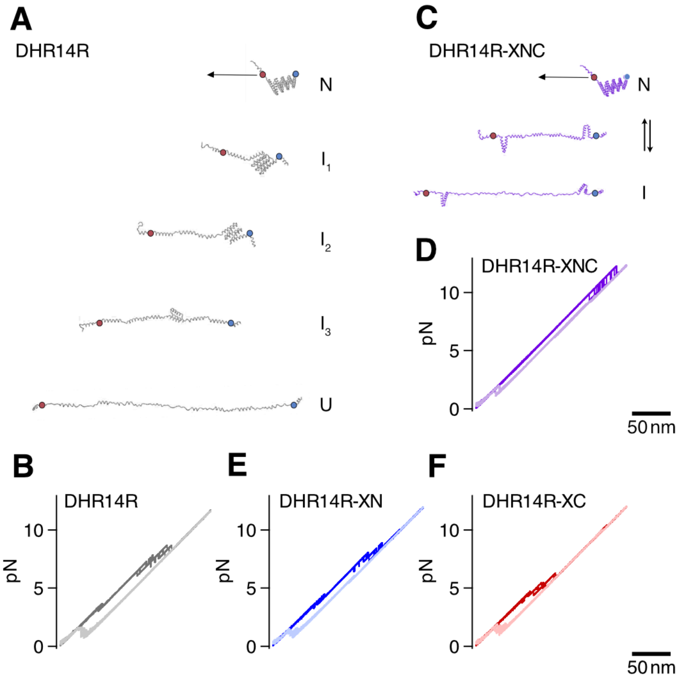

FIG. S2. **(A)** Structures of DHR14R intermediate states during (un)folding found by coarse-grained molecular dynamics simulations. Exemplary snapshots show that DHR14R unfolds from the native state (N) via three intermediate states ($I_1$–$I_3$) until the fully unfolded state (U) is reached. The N- and C-terminal pulling points are highlighted in blue and red circles, respectively. **(B)** Structures of double crosslinked DHR14R-XNC (un)folding states found by coarse-grained molecular dynamics simulations. DHR14R-XNC unfolds mainly in a single transition from the native state (N) to an intermediate state (I) in which the core is unfolded while the terminal crosslinked repeats remain structured. The N- and C-terminal pulling points are highlighted in blue and red circles, respectively. **(D–F)** Overlay of three independent simulated unfolding (dark shades) and refolding trajectories (light shades) for DHR14R (gray), DHR14R-XNC (purple), DHR14R-XN (blue), and DHR14R-XC (red). Consistent with experimental data, simulations for DHR14R showed unfolding and refolding via three intermediate states, corresponding to the unfolding of a single terminal helix, followed by the subsequent unfolding of two additional helices at a time. The last transition corresponded to the (un)folding of three helices, in agreement with experimental data. DHR14R-XNC showed mainly a single transition from the fully folded state (N) to the unfolded core, while the termini remained folded. Thus, the number and macroscopic conformation of the intermediate states observed in CG unfolding and refolding simulations are consistent with experimentally determined intermediate states.

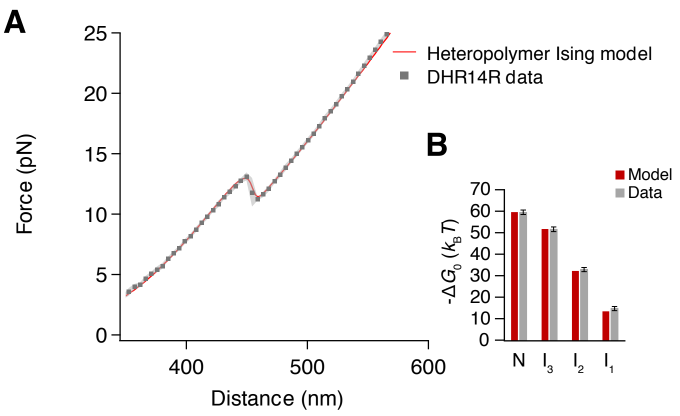

FIG. S3. Fitting a hetero-polymer Ising model to force-distance curves of DHR14R unfolding. **(A)** Fitted hetero-polymer Ising model (red curve) to measured Force-distance curves for DHR14R according to Synakewicz et al. (2022) [8]. The average for 96 FDCs is shown as gray dots with SD (shaded region). DNA fit parameters were adapted from the fitted experiment with K of 1000 pN/nm, $L_{\mathrm{DNA}}$ of 370 nm, $p_{\mathrm{DNA}}$ of 16 nm, combined trapstiffness of 0.32 pN/nm. **(B)** Energies of the modeled and measured DHR14R states. $\Delta G_0$ is the energy difference between the indicated states (N, $I_1$–$I_3$) and the unfolded state (U) based on the number of folded A and B helices in the corresponding conformation. Energies according to the Ising model in red and quantified energies from DHR14R force-distance measurements in gray. Error bars show standard deviations from $n = 9$ molecules. According to the Ising model, the intrinsic energy of a repeat $G_{\mathrm{unit}}$ is $0.5\,k_{\mathrm{B}}T$ and the coupling energy of neighboring repeats $G_{\mathrm{nn}}$ equals $-19\,k_{\mathrm{B}}T$ for $n = 4$ repeats. Additionally, we considered that the terminal repeats are intrinsically more stable due to solvent exposure, which is reflected in additional capping energies of $-1.5\,k_{\mathrm{B}}T$ for the N-terminal repeat and $-2.5\,k_{\mathrm{B}}T$ for the C-terminal repeat. Thus, we expect a weak stability of the capping repeats at room temperature (MD simulations). Energy details are listed in table S1

.

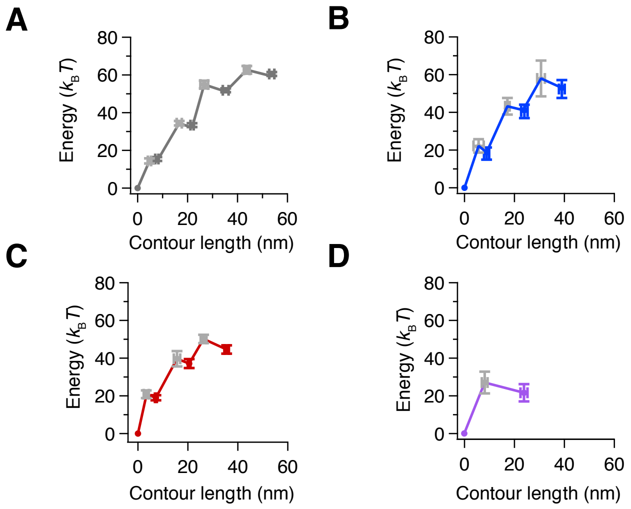

FIG. S4. **(A)** Corresponding interpolated energy landscape of DHR14R. Transition state positions are depicted in light gray. The barrier height was estimated using the Arrhenius equation. Error bars show statistical errors (SD) from $n = 9$ molecules. **(B-D)** Corresponding interpolated energy landscape of DHR14R-XN, DHR14R-XC, and DHR14R-XNC, n=6 molecules per construct.

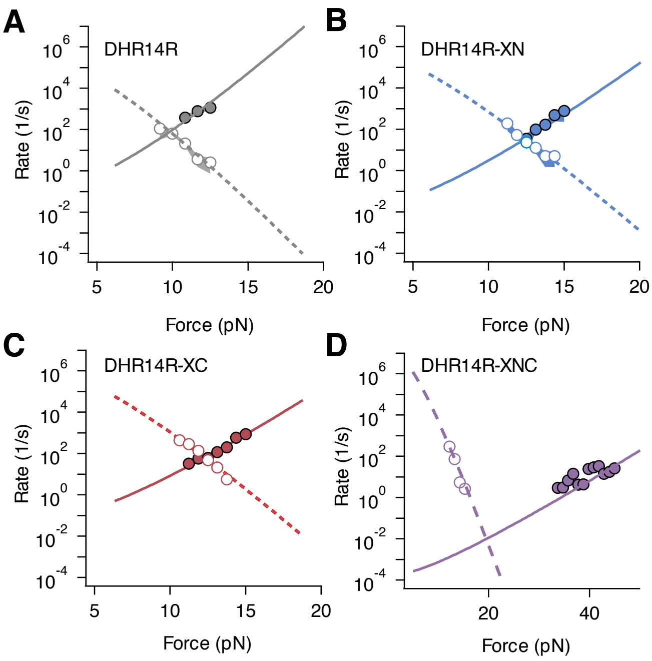

FIG. S5. Representative chevron plots for the first folding transition of all DHR14R variants. **(A)** DHR14R WT, DHR14R-XN **(B)**, DHR14R-XC **(C)**, DHR14R-XNC **(D)** respectively. Filled circles show unfolding, while empty circles represent folding events. Data were obtained with constant velocity pulling and relaxing cycles at 100 nm/s, except for DHR14R-XNC, for which a velocity of 500 nm/s was required to obtain sufficient data points for proper fitting.

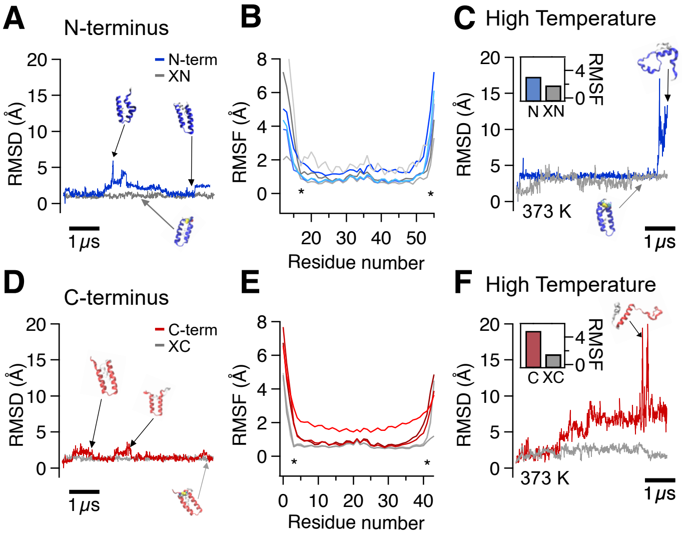

FIG. S6. All-atom molecular dynamics simulations of DHR14R N- and C-terminus in the absence and presence of a crosslink. **(A)** Example root-mean-square deviation (RMSD) trajectory from MD simulations for 5 μs at 298 K of DHR14R WT N-terminus in the absence (blue) and presence (gray) of a crosslink. **(B)** Root-mean-square fluctuation (RMSF) from MD simulations in triplicate without crosslink (blue shades) and with crosslinks (gray shades). The crosslinked residues are marked with asterisks. **(C)** RMSD trajectories for non-crosslinked (blue) compared to crosslinked (gray) N-terminal repeat simulated at 373 K.**(D)** Exemplary RMSD trajectories for the C-terminal repeat without (red) and with crosslink (gray), respectively. **(E)** RMSFs in triplicate for C-terminal repeat. **(F)** RMSD trajectories for non-crosslinked (red) compared to crosslinked (gray) C-terminal repeat simulated at 373 K

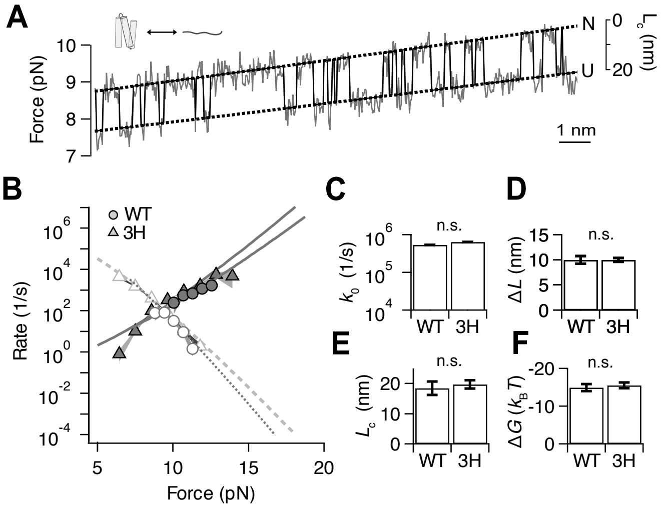

FIG. S7. **(A)** (Un)folding trace of the three C-terminal helices from DHR14R between the native (N) and the unfolded (U) state. **(B)** Exemplary chevron plot of folding/ unfolding kinetics for the full-length DHR14R (circles, called WT) compared to the truncated 3-Helix bundle (triangles, named 3H). **(C–F)** Fit parameters for the full-length WT compared to the truncated 3-Helix bundle (3H), $n$ = 6 molecules for DHR14R-3H. Error bars show statistical errors (SD).

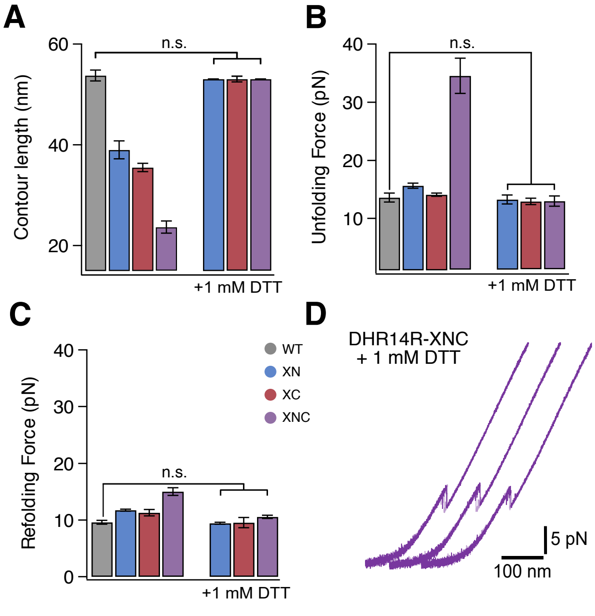

FIG. S8. **(A)** Change in contour length **(B)** Unfolding forces **(C)** Refolding forces, $n = 3$ molecules for reduced variants. Error bars show statistical errors (SD). **(D)** Exemplary FD cycles for reduced DHR14R-XNC construct. Stretch (dark shades) and relax (light shades) cycles were performed at 100 nm/s. For all constructs, the unfolding and refolding forces, the change in contour length, and the free energy of the WT DHR14R without any cysteines were reproduced within errors. Thus, the observed changes in mechanical stability and folding kinetics were exclusively caused by the introduced intramolecular crosslinks. No significant effect of substituting the selected residues with cysteine is observed.

#### IV. SUPPLEMENTARY TABLES

| $G_\text{A}$ | $G_\text{B}$ | $G_\text{AB}$ | $G_\text{AA}$ | $G_\text{BB}$ | $G_\text{BA}$ | $G_\text{N-cap}$ | $G_\text{C-cap}$ | $G_\text{unit}$ | $G_\text{nn}$ |
|---|---|---|---|---|---|---|---|---|---|
| 8.5 | 6 | −14 | −1.5 | −2 | −15.5 | −1.5 | −2.5 | 0.5 | −19 |

TABLE S1. Energy contributions in DHR14R model by the described heteropolymer Ising model. $G_\text{A}$: intrinsic energy of a type A helix, $G_\text{B}$: intrinsic energy of a type B helix, their coupling energy $G_\text{AB}$, $G_\text{AA}$, $G_\text{BB}$, and $G_\text{BA}$ are coupling energies of the respective helices in neighboring repeats, ($G_\text{N-cap}$): additional energy of the N-terminal cap, ($G_\text{C-cap}$): additional energy of the C-terminal cap. All energies are in $k_\text{B}T$.

#### V. SUPPLEMENTARY VIDEOS

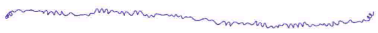

Video S1. Coarse-grained unfolding simulation DHR14R WT

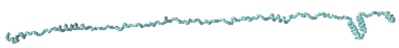

Video S2. Coarse-grained unfolding simulation DHR14R-XN with a N-terminal crosslink

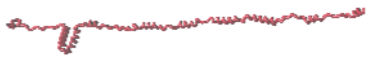

Video S3. Coarse-grained unfolding simulation DHR14R-XC with a C-terminal crosslink

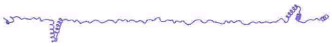

Video S4. Coarse-grained unfolding simulation DHR14R-XNC with N- and C-terminal crosslinks

[1] C. Bustamante, J. F. Marko, E. D. Siggia, and S. Smith. Entropic Elasticity of $\lambda$-Phage DNA. *Science*, 265(5178):1599–1600, September 1994.

[2] Marko, J. F and E. D. Siggia. Stretching DNA. 1995.
